# Frequency-dependent and capacitive tissue electrical properties in spinal cord stimulation models

**DOI:** 10.1101/2024.11.22.624883

**Authors:** Niranjan Khadka, Boshuo Wang, Marom Bikson

## Abstract

**Background:** Models of spinal cord stimulation (SCS) simulate the electric fields (E-fields) generated in targeted tissues, which in turn can predict physiological and then behavioral outcomes. Notwithstanding increasing sophistication and use in optimizing therapy, SCS models typically calculate E-fields using the quasi-static approximation (QSA). QSA, as implemented in neuromodulation models, neglects the frequency dispersion of tissue conductivity, as well as propagation, capacitive, and inductive effects on the E-field. The reliability of QSA specifically for SCS has not been considered in detail, especially for higher-frequency SCS.

**Methods:** We implemented a frequency-dependent finite element method (FEM) and solved a high-resolution RADO-SCS model with voltage-controlled (VC) and current-controlled (CC) stimulation to assess the impact of frequency-dependent conductivity (dispersion) and permittivity of spinal tissues on E-fields generated at three different spinal column locations (epidural space, spinal cord, and root) for frequencies spanning from 1 Hz to 10 MHz. Results were compared with predictions of QSA method, with varied conductivity values of purely resistive tissues. We further assessed the impact of frequency-dependent and capacitive tissue properties on spinal heating and distortion of the E-field waveform.

**Results:** Tissue-specific electric properties around the energized leads and mode of stimulation-control impacted the magnitude of E-fields. In the spinal cord, the VC-SCS E-field generated with the frequency-dependent and capacitive properties was comparable to the QSA with 2X epidural fat conductivity, whereas the CC-SCS generated E-field was minimally impacted by frequency-dependent and capacitive properties up to 10 kHz. Spinal cord heating predicted by frequency-dependent and capacitive tissue properties was comparable to the QSA conditions with VC-SCS, whereas with CC-SCS, there was no impact of the frequency-dependent and capacitive tissue properties in spinal cord heating. E-field waveform distortion in the spinal cord, with CC-SCS at 1 kHz-specific electrical properties, was significant when fat capacitance (permittivity) was increased by 10X, whereas with VC-SCS, there was no effect of tissue capacitance.

**Conclusion:** Regardless of the mode of SCS, QSA was still valid in predicting SCS-induced E-field and heating at the spinal tissues- across *α* and *β* dispersion region of spinal tissue’s dielectric spectrum for VC-SCS and up to 10 kHz for CC-SCS.

## Introduction

Computational modeling of neuromodulation current flow applies the quasi-static approximation (QSA)^1^: (1) no wave propagation or self-induction in tissue, (2) linear tissue properties, (3) purely resistive tissue, and (4) non-dispersive tissue. The purely resistive simplification is only valid at lower frequencies where resistive current dominates tissue electrical behavior, whereas at higher frequencies, the capacitive effects become more significant ^2–4^. The QSA is generally considered valid for frequencies below 10 kHz in biological tissues, with higher frequencies requiring consideration of complex tissue permittivity ^1,5–7^. Considering the frequency-dependent electrical properties can result in more accurate model predictions of stimulation waveforms and neural activation patterns compared to quasi-static models, especially for certain stimulation waveforms and frequency ranges.

The dose of spinal cord stimulation (SCS), i.e., the applied electric field (E-field) to the neural targets in the cord, depends on electrode polarity, waveform shape (frequency, pulse width, and amplitude), and complex electrical properties of the spinal tissues. Previously, through phantom studies and QSA computational modeling, we considered the role of purely resistive spinal tissues in electric field (E-field) generated during 10 kHz-SCS and HD-SCS ^8–10^. Here, to understand the role of frequency-dependent complex tissue electrical properties, we implemented the prior RADO-SCS model ^11^ and predicted E-fields generated by voltage-controlled SCS (VC-SCS) and current-controlled SCS (CC-SCS) at representative frequencies across the spectrum.

## Methods

We adapted a previously developed high-resolution RADO-SCS model ^11^ to estimate the role of frequency-dependent spinal tissue electrical properties in E-fields generated by clinical SCS intensities (VC-SCS and CC-SCS). Separately, the extent of spinal tissue local heating by SCS E-fields was also estimated. E-field and temperature predicted by frequency-dependent finite element method (FEM) was compared with QSA FEM method. For QSA, spinal tissue compartment conductivities were based on Gabriel et al., 1996 at 10 Hz^12^. The relative importance of not relaxing the purely-resistive tissue assumption of QSA was considered by varying spinal tissue conductivity under QSA conditions and compared with conditions with tissue capacitance included (Figure 3, Figure 4). The impact of tissue capacitance on induced E-field for a pulsed waveform at 1 kHz was also considered by comparing the baseline spinal tissues permittivity condition with a 10× change in epidural fat and spinal cord permittivity conditions, for both VC- SCS and CC-SCS.

### Frequency-dependent conductivity and permittivity

The tissue compartment-specific bioelectrical properties (relative permittivity (*ε*_r_) and electrical conductivity (*σ*)) as a function of frequency (*f*) were obtained from prior literature ^9,12,13^ and illustrated in Figure 1. The frequency range covers conventional and high-frequency SCS as well as diathermy applications ^8,9,14–17^, and includes the *α*- and *β*-dispersion regions of tissue’s dielectric spectrum^12^. The relative permittivity of biological tissues generally decreases with increasing frequency, except for some non-vascular tissues such as cerebrospinal fluid (CSF) and intervertebral disc, whereas the electrical conductivity was relatively constant across the spectrum (Figure 1). In addition, the capacitive conductivity (*ωε*, with the absolute tissue permittivity *ε* = *ε*_r_*ε*_0_, the permittivity of free space *ε*_0_=8.85×10^−12^ F·m^−1^, and the angular frequency *ω* = 2π*f*) and the capacitive-to-resistive conductivity ratio (*ωε*/*σ*) of biological tissues generally both increase with frequency.

**Figure 1:**
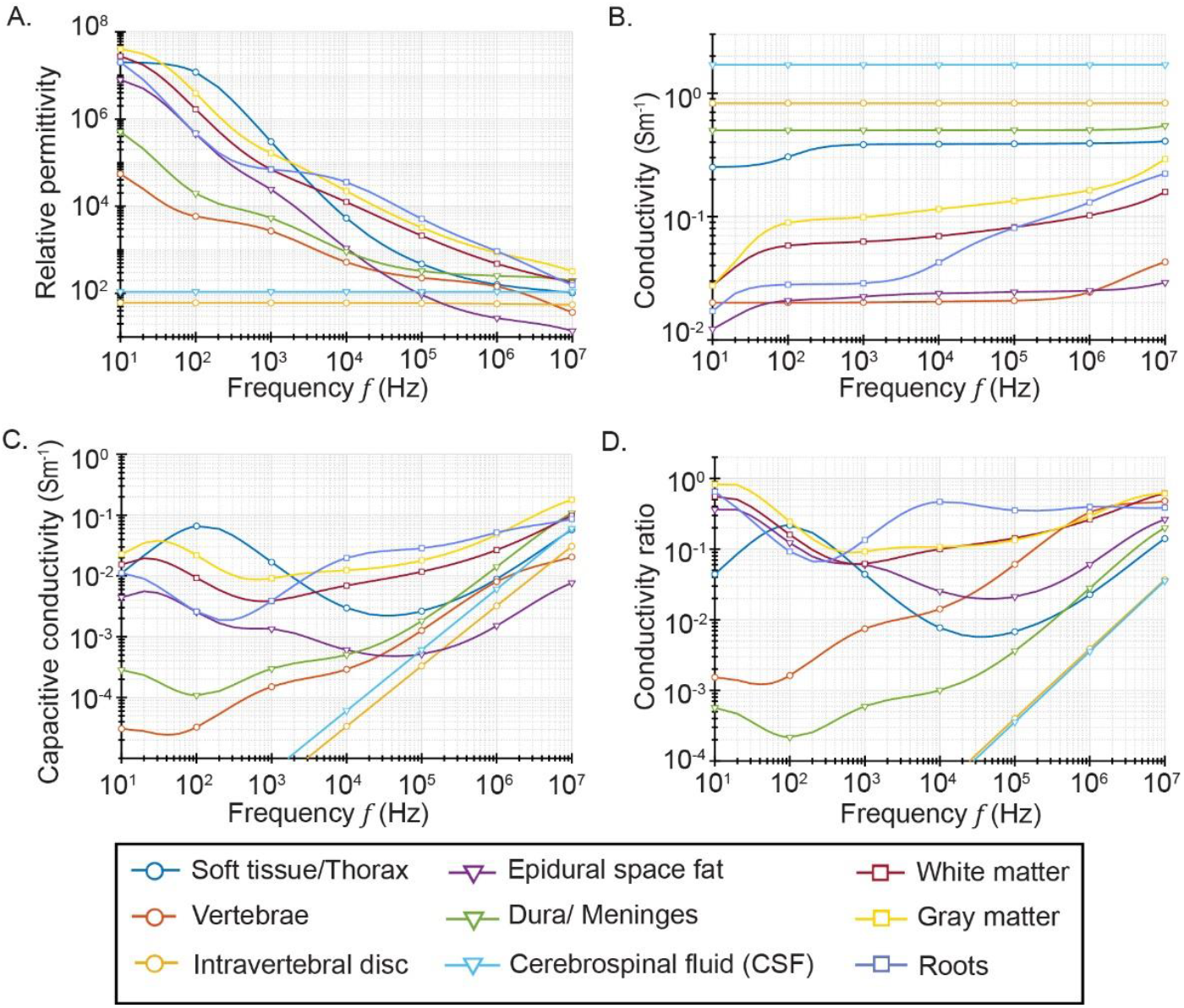
Relationship between simulated bioelectrical properties of spinal tissues at different frequencies. (A) shows relationship of relative permittivity (*ε*_*r*_), (B) electrical conductivity (*σ*), (C) capacitive conductivity (*ωε*), and (D) resistive-to-capacitive conductivity ratio (*ωε*/*σ*).

### Electric currents and fields

When tissue capacitance is considered, the total impressed (source) electric current density (***J***_t_) is related to the E-field (***E***) in the frequency domain by:

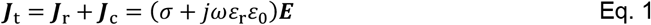

where ***J***_r_ and ***J***_c_ are the resistive and capacitive current density, respectively. The voltage distribution produced by an electrical stimulation during SCS in each spinal tissues with homogenous frequency-dependent electrical properties is solved according to conservation of current:

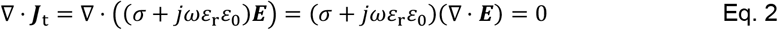

Hence, for conservative field that be expressed as the gradient of the induced voltage ***E*** = −Δ*V*, the Laplace’s equation holds within each tissue domain:

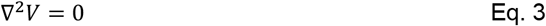

### Implementation of frequency-dependent electrical properties in SCS model

A state-of-the-art RADO-SCS ^11^ model with eight anatomically precise tissue compartments including, the lower thoracic region vertebrae (T8–T11), intervertebral disc (IV Disc), epidural fat, dura (meninges), CSF, white-matter, gray matter, and roots were included in the final assembly (Figure 2). We modeled an eight-contact clinical SCS lead (electrode diameter: 1.35 mm, electrode contact length: 3 mm, inter-electrode insulation length: 1 mm) using Solidworks (Dassault Systemes Corp., MA, USA) and positioned it 1 mm distal to the mediolateral dorsal column midline at the T10 spinal level. The assembled CAD model was imported into Simpleware ScanIP (Synopsys Inc., CA, USA) to generate a FEM mesh. An adaptive tetrahedral refined mesh was generated using a built-in voxel-based meshing algorithm in Simpleware ScanIP. The resulting SCS model comprised 70 million tetrahedral elements. The volumetric mesh was later imported into COMSOL Multiphysics 6.1 (COMSOL Inc., MA, USA) to solve the model using Laplace’s equation for E-field and Pennes’ bioheat equation for local heating, with a frequency-dependent solution method.

**Figure 2:**
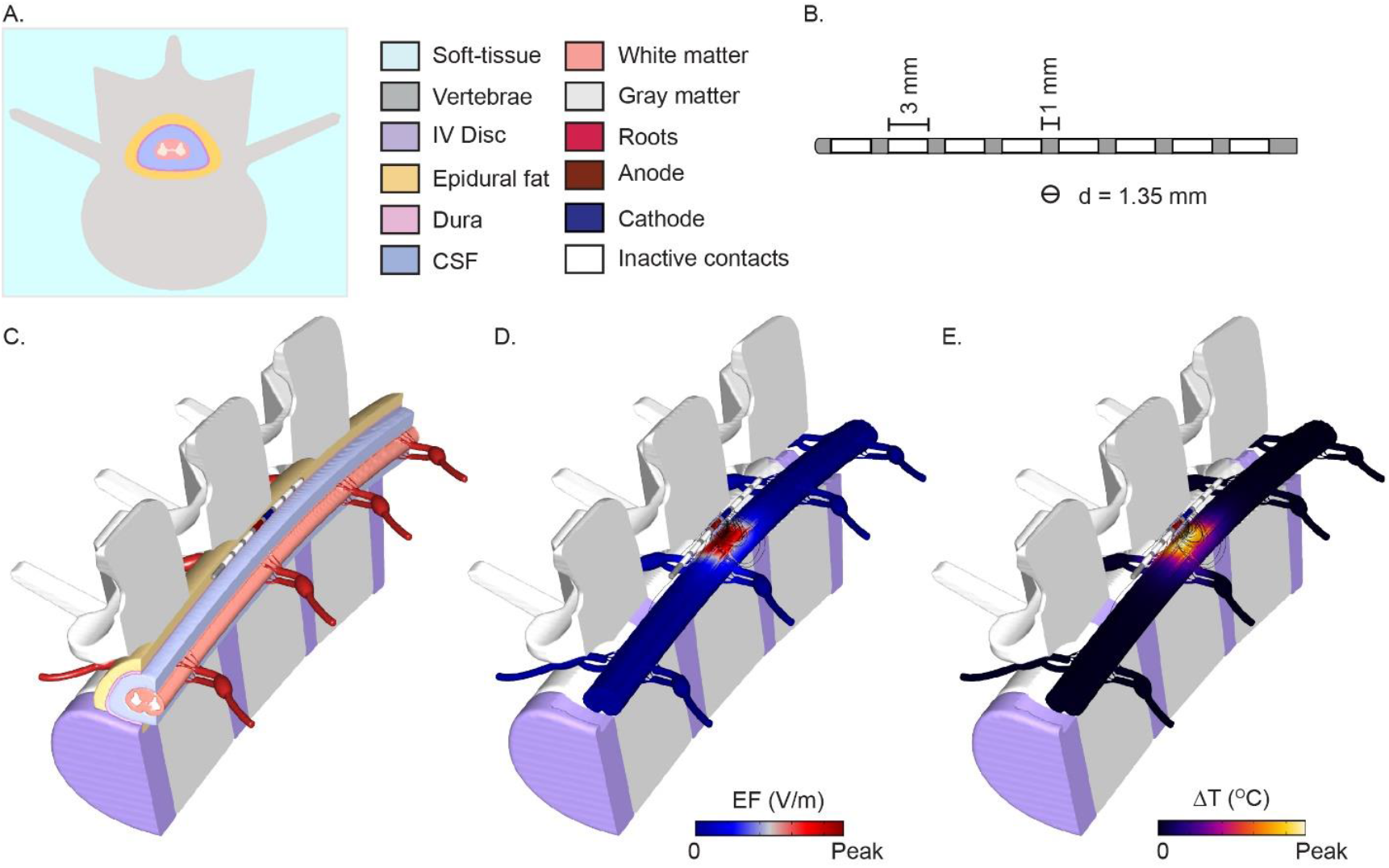
Anatomically detailed RADO-SCS model architecture and the predicted E-field and the temperature increase. (A) Color-coded anatomical compartments of the RADO-SCS model. (B) A clinical SCS lead and (C) a cut-out view of the 3D RADO-SCS model with the energized SCS lead contacts E5 (anode) and E4 (cathode) positioned 1 mm distal to the mediolateral dorsal column midline at the T10 spinal level. For an exemplary 3.5 mA_peak_ clinical SCS dose, (D) shows predicted E-field magnitude and (E) shows predicted temperature increase for a baseline spinal tissue conductivity and permittivity at 10 kHz.

For the QSA solution, a constant voltage/current was applied, via the anode while grounding the cathode. Frequency-dependent solution analyzes the SCS model response to varying frequencies, capturing dynamic behaviors (time-harmonic amplitude and phase), and is different from the QSA solution ^18–20^. To implement the frequency-dependent study in COMSOL, the applied current/voltage was specified as a variable rather than a constant using a ‘*Terminal’* boundary condition of type ‘*current/voltage*’, which allows for dynamic adjustments based on the model’s response to frequency of applied stimulus and solved using iterative solvers.

### Tissue heating in SCS model

The role of frequency-dependent electrical properties of spinal tissues in the extent of tissue heating by SCS was estimated by coupling the corresponding time domain solution of Eq. 1 (electrical stimulation) with the Pennes’ bioheat equation as:

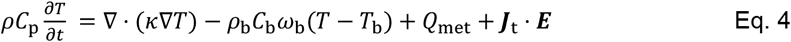

where *ρ, C*_p_, *T, T*_b_, κ, *ρ*_b_, *C*_b_, *ω*_b_, *Q*_met_, and ***J***_t_ · ***E*** represent tissue density (kg·m^−3^), specific heat (J·kg^−1^·K^−1^), temperature (K), blood temperature (K), thermal conductivity (W·m^−1^·K^−1^), blood density (kg·m^−3^), blood specific heat capacity (J·kg^−1^·K^−1^), blood perfusion rate (s^−1^), metabolic heat generation rate (W·m^−3^), and the differential form of the Joule heating due to SCS (W·m^−3^), respectively. On the slower timescale of bioheat phenomena, the Joule heating of sinusoidal E-field and current can be considered constant and replaced by the effective RMS value, 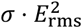,with the tissue capacitance not contributing to the heating. Considering the steady-state solution of tissue heating, the equation is solved with the time derivative of the temperature (∂*T*/∂*t*) set to zero.

The assigned thermo-biological properties of the biological tissues and thermo-electrical properties of the SCS lead are based on our previous studies ^9,11^. For the electrical boundary conditions, we energized the SCS lead contacts E5 (anode) and E4 (cathode) in a bipolar configuration using clinically-relevant SCS intensities (3.5 mA_peak_ and 3.5 V_peak_). The remaining external boundaries of the model were insulated, and continuity was assigned to the internal boundaries. Floating boundary conditions were also assigned to the remaining inactive electrodes in the model. For the thermal boundary conditions, the temperature at the outer boundaries of the spinal column was fixed at core body temperature (37°C) assuming no convective heat loss to the ambient temperature, no convective gradients across spinal surrounding tissues, and no SCS-induced heating at the model boundaries. The initial temperature of the tissues was set at 37°C. The relative tolerance was set to 10^−6^ to improve the solution accuracy. For both VC-SCS and CC-SCS, the resulting E-field distribution in the epidural space, spinal cord (white matter), and root (dorsal) with baseline tissue-specific permittivity was modeled and compared with the QSA solution (Figure 3, Figure 4). When indicated, the baseline permittivity of the epidural fat or spinal cord (both white matter and gray matter) were increased by 10× to estimate the role of tissue capacitance in E-fields generated in different spinal tissues and local tissue heating over different frequencies. For the QSA solution, the conductivity of the epidural fat or spinal cord were also increased or decreased by a factor of 2, to examine the sensitivity of the solution to these parameters^21^ and compare model prediction with the frequency-dependent solution.

**Figure 3:**
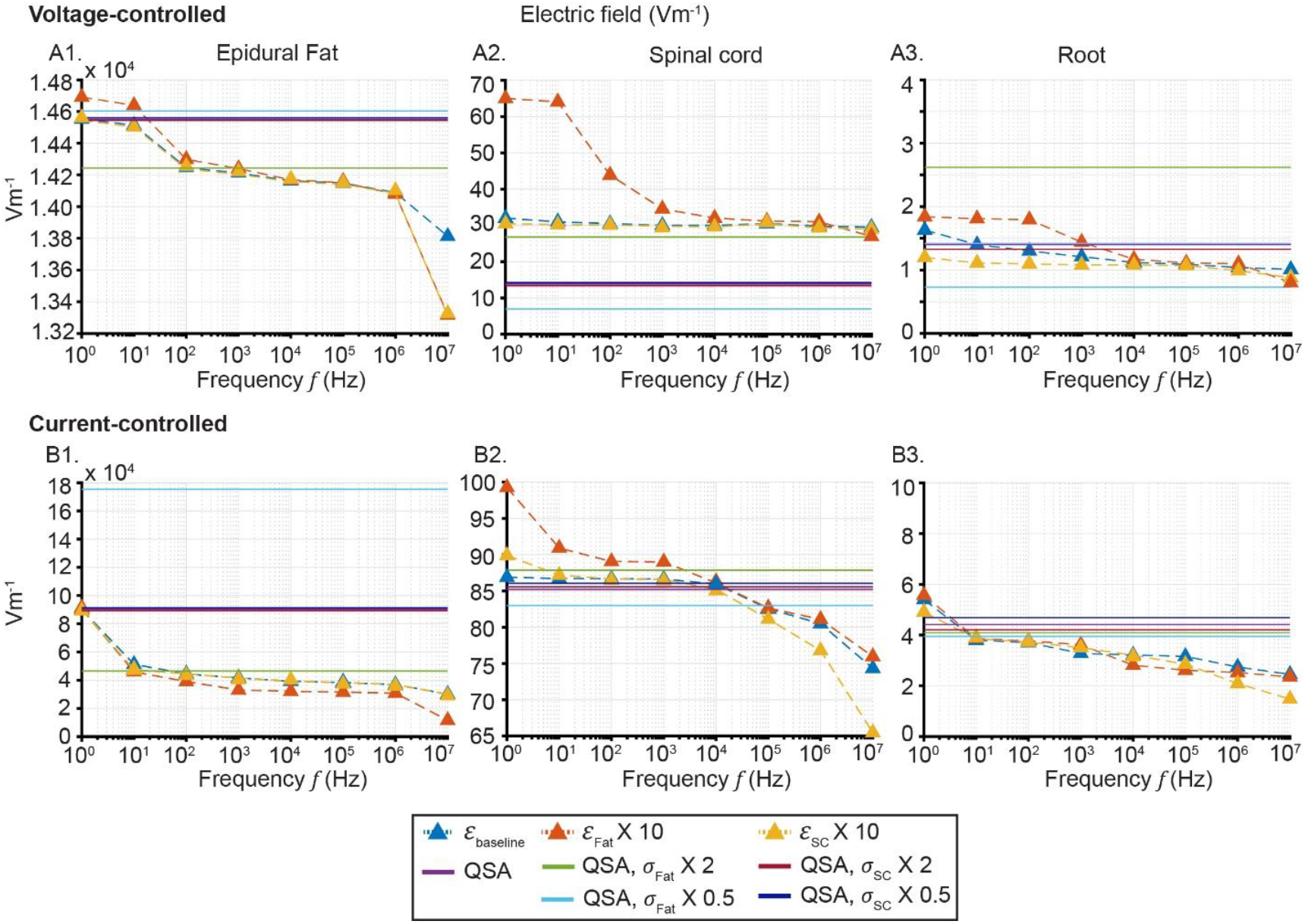
The peak E-field predicted in the epidural fat, spinal cord, and root for VC-SCS (A1, A2, and A3) and CC-SCS (B1, B2, and B3) with baseline spinal tissues permittivity, 10× epidural fat and spinal cord permittivity, and QSA conditions, across different frequencies. For VC-SCS, the predicted peak E-field in the spinal cord was comparable to QSA with 2X fat conductivity, whereas for CC-SCS, the predicted peak E-field in the spinal cord was minimally higher up to 10 kHz with frequency-dependent and capacitive tissue conditions compared to the QSA conditions.

**Figure 4:**
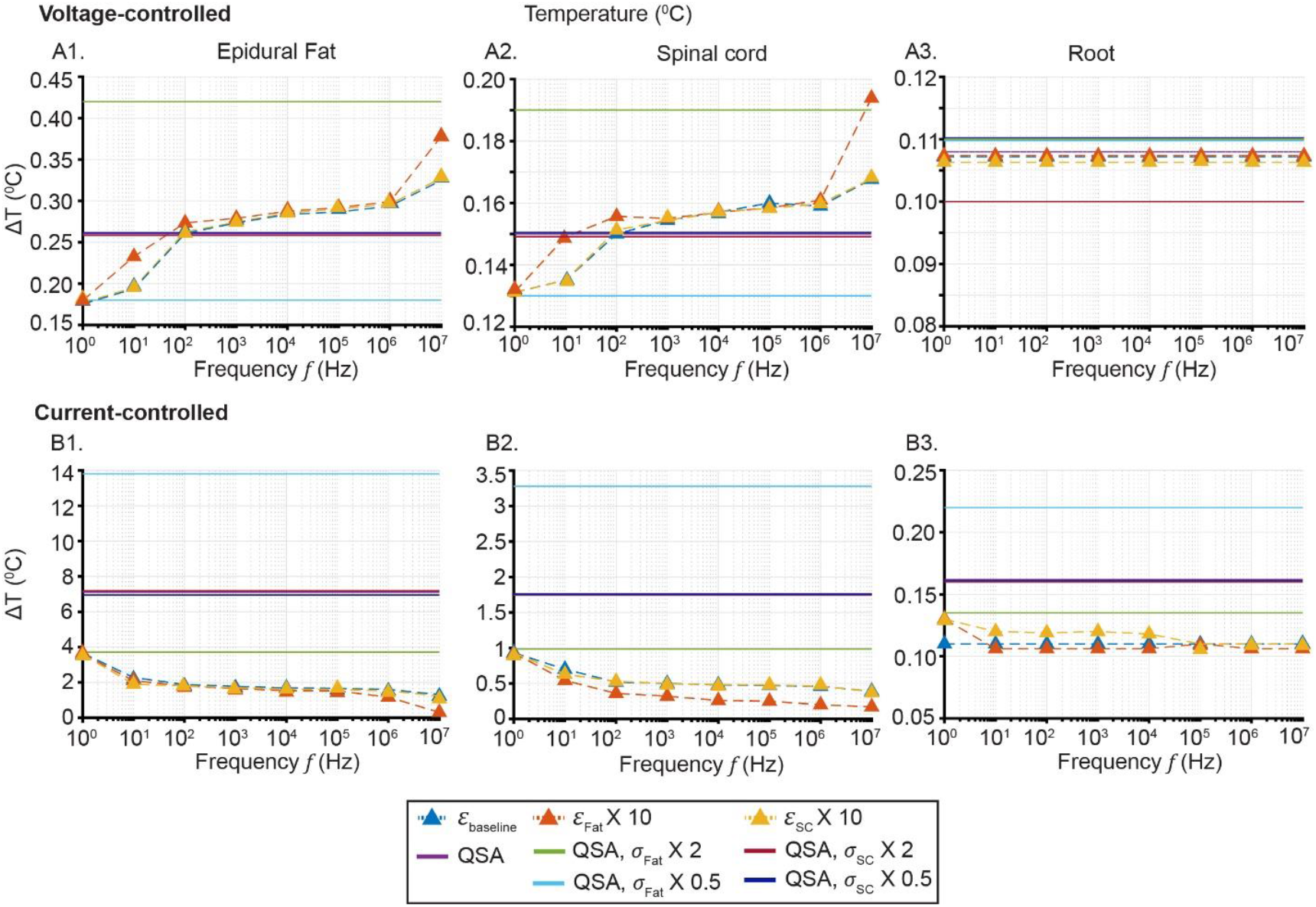
The peak temperature increase sampled in the epidural fat, spinal cord, and root for VC- SCS (A1, A2, and A3) and CC-SCS (B1, B2, and B3) with baseline spinal tissues permittivity, 10× epidural fat and spinal cord permittivity across frequencies, and QSA conditions. Temperature increase with frequency-dependent and capacitive property conditions was comparable for VC-SCS and lower for CC- SCS in all spinal tissue locations, compared with the QSA conditions.

### E-field waveform distortion due to tissue capacitance

In a separate time-dependent analysis (Figure 5), the role of frequency-dependent permittivity (from baseline permittivity at 1 kHz to 10× epidural fat and spinal cord permittivity) in E-field waveform distortion was also assessed. We applied a monophasic pulse of 400 µs pulse width, 900 µs pulse duration, and 3.5 V_peak_ (VC-SCS) and 3.5 mA_peak_ (CC-SCS) intensity through contact E5 of the clinical SCS lead, with contact E4 as ground/return. The distortion in the generated E-field waveform was assessed for both modes of stimulation in the epidural fat and the spinal cord proximal to E5 for baseline permittivity condition and for a 10× increase in permittivity of the epidural fat or spinal cord from the baseline value.

**Figure 5:**
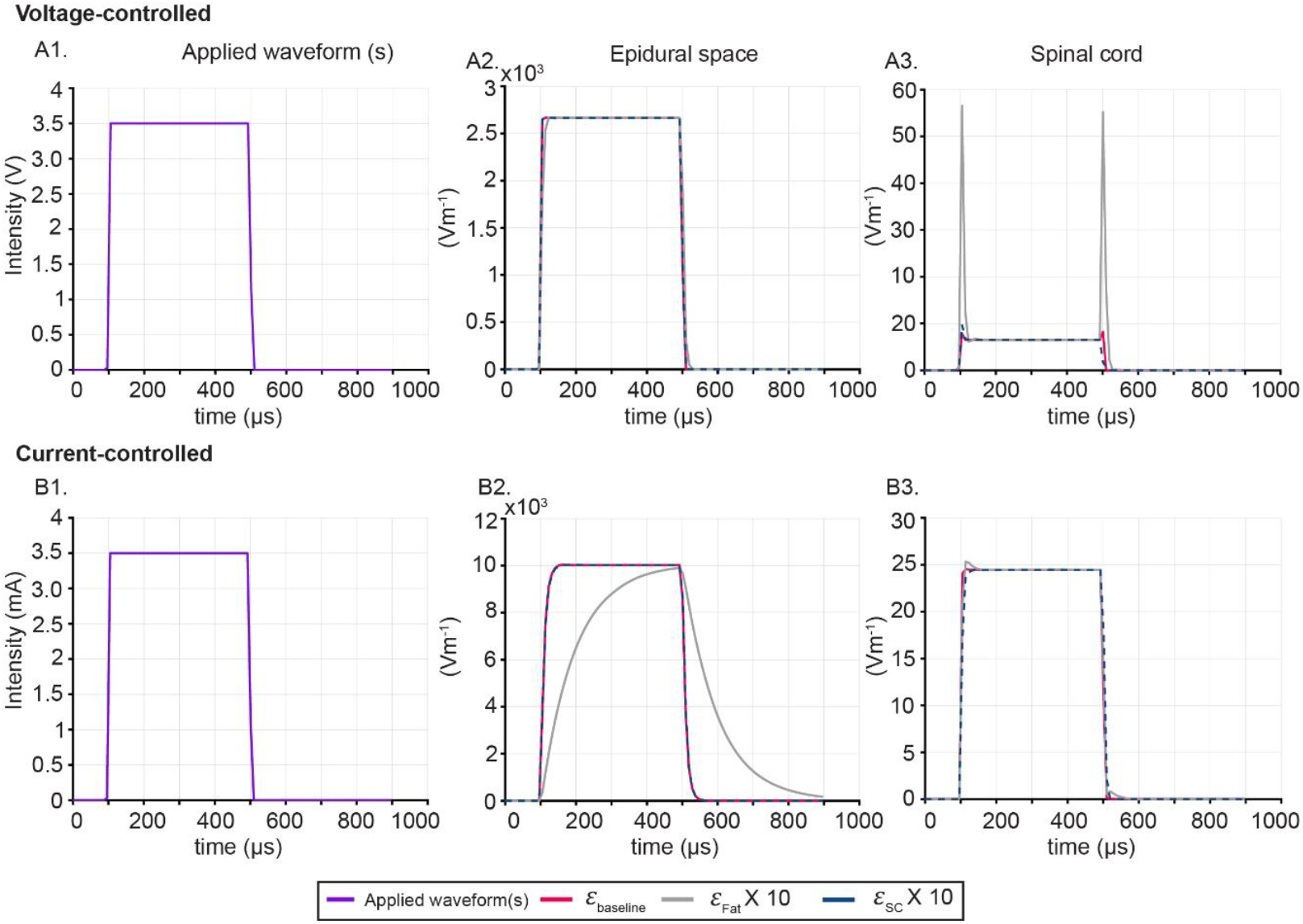
Role of spinal tissue permittivity in E-field waveform distortion in the epidural space and spinal cord for a VC-SCS (A1, A2, and A3) and a CC-SCS (B1, B2, and B3) with monophasic rectangular pulse of 400 µs duration and 3.5 V and 3.5 mA amplitudes, respectively applied via contact E5 of a clinical SCS lead. In the spinal cord, a 10X increase in epidural fat permittivity predicted distorted E-field waveform with spikes at pulse edges for VC-SCS, whereas for CC- SCS, there was no effect of permittivity in E-field waveform distortion.

## Results

We implemented an anatomically realistic RADO-SCS model to assess the role of frequency-dependent and capacitive tissue properties of spinal tissues, considering either VC-SCS or CC- SCS, on generated E-field and tissue heating. The following tissue conditions were compared: 1) ‘baseline’ of frequency-dependent and capacitive tissue properties; 2) frequency-dependent properties with 10 fold increases of the permittivity of fat or spinal cord (to access the specific impact of tissue capacitance); 3) QSA using either ‘standard’ tissue resistivities or with doubling or halving of the resistivity of fat or spinal cord. The variation (0.5x or 2x) of tissue resistivity in the QSA model provided context for the relative importance of frequency-dependent and capacitive tissue properties. Peak E-field in each tissue compartment was predicted as a function of applied stimulation frequency content at 3.5 V_peak_ (Figure 3A) for VC-SCS and 3.5 mA_peak_ for CC-SCS (Figure 3B). All results are reported at 3.5 V_peak_, or 3.5 mA_peak_, but E-field scale linearly with applied intensity so that relative effects at any intensity can be inferred.

With limited exceptions, there was a minimal impact of frequency-dependent and capacitive properties in peak E-field at any stimulation mode, location, or frequency (Figure *3*). Spinal cord heating predicted by frequency-dependent and capacitive tissue properties was comparable to QSA conditions with VC-SCS, whereas with CC-SCS, there was no impact of frequency-dependent and capacitive tissue properties in spinal cord heating (Figure *4*). E-field waveform distortion in the spinal cord, with CC-SCS at 1 kHz-specific electrical properties was only significant when fat capacitance (permittivity) was increased by 10X, whereas with VC-SCS, there was no effect of tissue capacitance (Figure *5*).

## Discussion and conclusion

Prior SCS models utilized QSA and only assumed fixed (frequency independent) resistive properties of the biological tissues with no tissue capacitance. However, biological tissue exhibits frequency-dependent electrical properties ^3,13,22–24^ that will change the SCS generated E-fields compared to QSA solutions. In this computational modeling study, we considered the frequency-dependent electrical property of the spinal tissues at VC-SCS and CC-SCS to estimate the peak E-field at different frequencies and compared with QSA conditions. Electrode capacitance, excluded from this study, significantly influences E-field transmitted to the spinal tissues during stimulation ^25,26^. Both VC-SCS and CC-SCS models tested clinically-used intensities. Though the predicted E-fields were different for the two modes of stimulation (different intensities to start with—3.5 mA_peak_ versus 3.5 V_peak_), they both showed dependence on the permittivity of spinal tissues.

We assessed the sensitivity of predicted E-fields of VC-SCS and CC-SCS to changes in tissue electrical properties across different frequencies, however, we did not explicitly model E-field distortion with varying frequency of stimulation. For both VC-SCS and CC-SCS, the QSA conditions predicted comparable E-field in the spinal cord as the frequency-dependent and capacitive tissue property conditions suggesting validity of the QSA for SCS modeling. Tissue heating predicted for both modes of SCS with frequency-dependent and capacitive property conditions didn’t exceed the QSA conditions.

## Notes

**Conflict of Interest** The City University of New York holds patents on brain stimulation with MB and NK as inventors. NK consults for Ceragem Medical. MB has equity in Soterix Medical Inc. MB consults, provides expert witness support, received grants, assigned inventions, and/or served on the SAB of SafeToddles, Zabara Family Foundation, Boston Scientific, GlaxoSmithKline, Biovisics, Axonics, Mecta, Lumenis, Halo Neuroscience, Wave Neuroscience, Google-X, i-Lumen, Humm, Allergan (Abbvie), Apple, Ybrain, Ceragem, Ceragem Clinical, Remz. MB is supported by grants from Harold Shames and the National Institutes of Health: NIH-NIDA UG3DA048502, NIH-NIGMS T34 GM137858, NIH-NINDS R01 NS112996, NIH-NINDS R01 NS101362, and NIH-G-RISE T32GM136499.

### Competing Interest Statement

The City University of New York holds patents on brain stimulation with MB and NK as inventors. NK consults for Ceragem Medical. MB has equity in Soterix Medical Inc. MB consults, provides expert witness support, received grants, assigned inventions, and/or served on the SAB of SafeToddles, Zabara Family Foundation, Boston Scientific, GlaxoSmithKline, Biovisics, Axonics, Mecta, Lumenis, Halo Neuroscience, Wave Neuroscience, Google-X, i-Lumen, Humm, Allergan (Abbvie), Apple, Ybrain, Ceragem, Ceragem Clinical, Remz. MB is supported by grants from Harold Shames and the National Institutes of Health: NIH-NIDA UG3DA048502, NIH-NIGMS T34 GM137858, NIH-NINDS R01 NS112996, NIH-NINDS R01 NS101362, and NIH-G-RISE T32GM136499.

